# Life history trade-offs in glucocorticoid-mediated habitat selection

**DOI:** 10.1101/2022.03.03.482820

**Authors:** Levi Newediuk, Gabriela F. Mastromonaco, Vander Wal Eric

**Affiliations:** Department of Biology, Memorial University, St. John’s Newfoundland, Canada A1B 3X9; Reproductive Physiology, Toronto Zoo, Toronto Ontario, Canada M1B 5K7

## Abstract

The cort-adaptation and cort-fitness hypotheses both propose glucocorticoids produced by parents mediate trade-offs between their own survival and that of their offspring. The contribution of glucocorticoids to offspring provisioning can quantify these trade-offs because provisioning poses a risk to parents. However, attributing provisioning behaviour to glucocorticoids is difficult because glucocorticoids often drive foraging behaviours to store energy for later provisioning. We compared the effects of glucocorticoids on habitat selection before and after calving to test for the trade-offs female elk make when provisioning offspring. Despite finding that female elk with elevated glucocorticoids selected more strongly for high-risk, high-forage cropland, we found no difference in glucocorticoid production nor a significant change in the effect of glucocorticoids on cropland selection at calving time when lactating elk required the more energy. However, we found a gradual increase in the effect of glucocorticoids on cropland selection by female elk as their calves grew, suggesting the growing energy requirements of calves encouraged more risky habitat selection behaviours by their parents over time. We suggest trade-offs in investment between parents and offspring across life history stages can be tested by integrating glucocorticoids into habitat selection models. Ultimately, this integration will help elucidate the adaptive function of glucocorticoids.

## Introduction

A key question in animal ecophysiology is whether glucocorticoids, the so-called “stress” hormones, drive behaviours that increase fitness or indicate exposure to stressors that might compromise fitness. Negative relationships often emerge between glucocorticoids and fitness because glucocorticoid production is one common response among vertebrates to dealing with stressful environments (Bonier et al., 2009a). Stress, however, is something all organisms have evolved to deal with, and there is much evidence that the conserved production of glucocorticoids among all vertebrates to deal with stressors is adaptive (Dantzer, 2023; Dantzer et al., 2013). Consistent with this adaptive theory of glucocorticoid production, two competing hypotheses predict that glucocorticoid production instead increases fitness by supporting the behaviours individuals use to continue to survive and reproduce in stressful environments. For example, the “cort-adaptation” hypothesis proposes glucocorticoids increase fitness in stressful environments because their production stimulates behaviours like foraging and provisioning to meet the energetic demands of reproduction (Bonier et al., 2009a). However, behaviours that support reproduction also increase risks to parents and challenge their own growth and survival. The alternative “cort trade-off” hypothesis instead theorizes glucocorticoids moderate investment in reproduction, stimulating behaviours like brood abandonment that support parent survival at the expense of offspring (Love et al., 2004).

Expectations of both the cort trade-off and cort-adaptation hypotheses are supported by the manifold behavioural effects of glucocorticoids. For example, one role of glucocorticoids is to support self-preservation behaviours like escape from predators. Escape behaviours require energy, and glucocorticoids mobilize energy by moving glucose from storage into the bloodstream (Dallman et al., 1995). Mobilizing energy for escape from predators is adaptive in most contexts. However, unless reproducing under severe stress that threatens their own survival, parents might compromise their fitness with escape during reproduction because it sometimes entails offspring abandonment (Love et al., 2004). Instead, glucocorticoid production by parents during reproduction often supports other adaptive behaviours. For example, glucocorticoids also feed back on the central nervous system with neurological effects, encouraging feeding behaviour (Dallman et al., 1995). This might be why many species actually produce their highest glucocorticoid levels of the year during reproduction when more foraging is required to provision offspring (Romero, 2002). However, while frequent provisioning supports reproduction, places with the best food resources also frequently pose the greatest risk of predation (Brown and Kotler, 2004), and thus provisioning also comes at a survival cost to parents (Ghalambor and Martin, 2001). Life history theory predicts that parents would only risk their own survival in stressful environments where they have fewer opportunities for future reproduction (Ghalambor and Martin, 2001). Thus, whether glucocorticoids drive provisioning or risk-taking behaviour during reproduction is needed to assess whether elevated glucocorticoids are evidence of stressful environments.

The primary challenge in understanding the fitness implications of glucocorticoids lies in attributing provisioning and risk-taking behaviour to glucocorticoid production. The association between risky provisioning and glucocorticoids is relatively straightforward in species like birds whose provisioning can be quantified by returning food to a nest. The more a bird parent visits a nest, the more food items it has acquired, and the more it has presumably risked its own survival to do so (Ghalambor and Martin, 2001). However, linking risky behaviours to glucocorticoids is comparatively complicated in other taxonomic groups. Mammals, for example, provision offspring with milk. Milk comes from food consumed and metabolized by the parent some time before provisioning offspring. The delay between consumption and provisioning means milk consumption is not necessarily proportional to the risks the parent has taken, sometimes over long periods of time, to build their energy stores for lactation. One of the ways both lactating mammals and birds avoid risk while acquiring energy is by regularly alternating between foraging habitat and refuge habitats with less forage (De Groeve et al., 2023; Dunning et al., 1992), the latter of which takes time away from foraging and thus eventual energy investment in offspring. As a result, a more universal representation of trade-offs between offspring and parent might be the overall proportion of time parents spend in safer refuge habitats with less forage, versus riskier foraging habitats where they acquire energy to invest some time later in offspring. Similarly, whether glucocorticoids support parents or offspring might be best determined by how glucocorticoids affect that habitat selection behaviour.

Habitat selection models are a well-established tool for assessing which variables affect habitat selection behaviour. Traditionally, these models measure how characteristics of habitats themselves influence which are selected or avoided (Boyce and McDonald, 1999). However, more recent interest in behavioural differences between individual animals has inspired innovative new models quantifying the effects of dynamic social environments (Webber et al., 2021), behavioural states (Picardi et al., 2022), and disease (Turner et al., 2021) on habitat selection. Another natural extension of habitat selection models in the context of ecophysiology would be to quantify how elevated glucocorticoid levels affect habitat selection during reproduction. We suggest integrating glucocorticoid levels into habitat selection models as a universal way of measuring the effects of glucocorticoids on trade-offs between parents and offspring, with the ultimate goal of distinguishing when glucocorticoids indicate stressful environments versus when they might increase fitness.

We used habitat selection models to test the role of glucocorticoids in driving the behaviour of female elk (*Cervus canadensis*) during reproduction. To reproduce, female elk must both maintain body condition prior to calving and compensate for the energetic demands of lactation (Cook et al., 2013). Like other species, these energetic demands are met by focusing foraging in areas with better forage that carry higher risk. Risk and forage quality in our study population are both highest in open cropland, and both diminish in safer habitats like forest and shrubland that make up the rest of the surrounding landscape (Hinton et al., 2020). We first predicted that elk would select for locations further from forest and shrubland — and thus closer to the better-quality forage in cropland — after they produced more glucocorticoids. Despite being risky, cropland provides the highest quality forage for elk in our study system (Hinton et al., 2020), whose energy requirements increase several-fold between the last day of gestation and their first day of investing in lactation (Nelson and Leege, 1982). We hypothesized that glucocorticoid production is the mechanism by which elk start to invest more in their offspring after reproduction. We predicted this change in investment could occur in two non-mutually exclusive ways. First, higher levels of glucocorticoids produced after calving might increase investment in finding better quality forage. Thus, we predicted mean glucocorticoid levels might increase after calving. Alternatively, the sensitivity of elk behavioural responses to glucocorticoids might change after calving without an overall change in glucocorticoid levels. Since provisioning calves by lactation is energetically expensive, we predicted glucocorticoids might cause even stronger selection for riskier habitats with better forage after calving. We also predicted the effect might become stronger with time post-calving to match the growing energetic requirements of the growing calf.

## Methods

### Estimating calving dates

We used global positioning system (GPS) locations of adult female elk to both characterize habitat selection and identify calving dates. In February 2019, 13 adult female elk were captured in southeast Manitoba, Canada (49.134, −96.557) from a population of approximately 150 individuals. Individuals were fit with GPS collars (Vertex Plus 830 g, VECTRONIC Aerospace GmbH, Berlin, Germany) that collected locations every 30 minutes during the calving season (May through July). We identified potential 2019 and 2020 calving sites by monitoring the mothers’ movement patterns. After locating calves, we fit each with a very high frequency (VHF) radio collar (V6C 83 g, Lotek, Newmarket, Ontario, Canada) for monitoring calf survival. Both adult female and calf capture procedures were in accordance with approved animal care protocols (Memorial University of Newfoundland animal use protocol #19-01-EV).

We processed all data and performed all statistical analyses using R v4.3.2 (R Core Team, 2023). We used location data from the adult female GPS collars to estimate unobserved calving events. Elk calves hide for 4-5 days following parturition until they are mobile enough to escape predators (Geist, 1982). This limited mobility causes elk mothers to reduce their own movement rates to remain close to the calf (Brook, 2010). We used the frequency of return visits to the potential calf to estimate parturition date using a machine learning approach (Marchand et al., 2021). We used the *recurse* package in R (Bracis et al., 2018) to calculate the number of return visits by each elk to within a buffer of each of its location points between May 15 and July 20 in both 2019 and 2020. Unlike some other ungulate species, elk calves select new hiding spots away from the calving site shortly after parturition (Johnson et al., 2006), meaning mothers might make return visits to different locations. To account for variation in return location, we used a 300 m radius buffer (Wallace and Krausman, 1992) to calculate recursive movements to the calf rather than the suggested 100 m radius buffer (Marchand et al., 2021).

We used elk movements surrounding 16 confirmed calving events as training data to predict an additional 11 unconfirmed events. We defined calving as the period between the confirmed calving date up to 5 d following to account for the most intensive hiding phase. After down sampling the training data to balance the number of points within and outside the calving period, we used a random forest classifier to predict the probability of each training data point belonging to the calving period. We averaged the probability of calving for each point falling within the known parturition period and used this as a threshold for detecting calving periods in the testing data. Specifically, we located where average probabilities exceeded the known calving threshold within a 5-d rolling window in the testing data. After repeating this process 100 times, we selected the 5-d window of points with the highest probability of belonging to the calving period for each calving event. We set the estimated the calving date as the first date within that period.

### Hormone sampling

We collected 181 fecal pellet samples to monitor glucocorticoid levels of the 13 collared elk from May–August 2019 and 2020. We identified clusters of location data indicative of bedding, and after confirming bedding by visiting the locations within 24 h of the individual being present in the area, we collected any visible fecal material. Because fecal glucocorticoid metabolites (FGM) are the product of circulating glucocorticoids metabolized over a period of hours to days (Gormally and Romero, 2020), elevated levels can indicate one or more responses to exogenous (e.g., predator encounters) or endogenous (e.g., calving) stressors during that time. For many ungulate species, FGM levels remain elevated for approximately 20 h following stress responses (Ashley et al., 2011; Escribano-Avila et al., 2013). The validated metabolization period for the Roosevelt elk (*Cervus canadensis roosevelti*) is 22 h (Ashley et al., 2011), and 18 h in red deer (*Cervus elaphus*), a close relative of the North American elk (Huber et al., 2003). This makes FGM an integrated proxy for both baseline and stress-induced circulating glucocorticoids during the 18-22 h preceding defecation.

Aside from individual differences in circulating levels, FGM levels are also influenced by environmental factors after defecation. These variables create discrepancies between circulating glucocorticoids and FGM that are not attributable to stressors. Specifically, FGM recovery from samples is affected by moisture in the field and failure to promptly store samples after collection (Romero and Wingfield, 2016). Thus, we avoided sampling after rain, collected samples within 24 hours of suspected defecation, and froze samples as soon as possible (< 8 hours) after collection (Sheriff et al., 2011).

We identified individuals by comparing DNA extracted from fecal samples to that from whole blood samples taken from individuals at the time of capture. However, like FGM concentration fecal DNA is susceptible to degradation from inclement weather and storage conditions, and only approximately 20% of extractions were successful. For those samples we could not identify using DNA (117 of 181 samples), we used supervised machine learning to assign suspected individuals to samples based on movement patterns and level of elk activity in the vicinity of the sample. The training model identified whether samples belonged to the suspected individual with 77% accuracy (Newediuk and Vander Wal, 2021; see also for further details on DNA extraction and machine learning models). We used this accuracy as a threshold for correct identification, predicting the accurate identification of testing samples over 500 iterations. We assumed samples belonged to the suspected individual when the mean predicted accuracy of testing samples exceeded the threshold accuracy. When mean predicted accuracy was less than the threshold, we tested whether samples could have belonged to a different collared individual in the same area around the time of defecation. We identified candidate individuals as those with any location points within 20 m of the sample up to 2 d prior to the time of sample collection. We repeated the same machine learning procedure for these new individuals, replacing the original individual that did not meet the threshold for correct identification with the new suspected individual. As above, we assumed samples belonged to the new individual if the predicted accuracy across 500 iterations exceeded the threshold accuracy.

### Statistical analysis

We tested for differences in glucocorticoid levels before and after calving using a Bayesian generalized linear model. Our model included the period before and after calving as a categorical variable and glucocorticoid levels in FGM samples as the response. We also included random intercepts for individuals to account for individual differences in glucocorticoid levels and scaled and centred glucocorticoid levels prior to analysis. We fit the model using the *brms* package in R (Bürkner, 2017), with a gaussian link function, weakly informative prior slopes with mean 0 and standard deviation 1, 4 chains, and 10,000 iterations including 5,000 warmup iterations.

To test whether glucocorticoid levels affected habitat selection, we used a version of habitat selection model called integrated step selection analysis (iSSA; Avgar et al., 2016). All habitat selection models, including iSSA, quantify the relative probability of selection for habitats using logistic regression, where the distribution of habitat values at used locations is compared to another sample of habitat values at available locations. Step selection analyses are a type of habitat selection analysis in which available locations are drawn from empirical distributions of step length and turn angles at each used location, thereby constraining available locations to the step level. However, habitat selection at each step depends on expected movement patterns. Without accounting for that movement in available step samples, model coefficients could be biased. To avoid this bias, iSSA steps are sampled from pre-specified distributions of turn angles and step lengths parameterized on observed steps (Avgar et al., 2016). Constraining available steps in this way simultaneously estimates movement and habitat selection coefficients (Fieberg et al., 2021), making it possible to test the effect of temporally variable factors like glucocorticoid levels on habitat selection throughout the movement process.

Our iSSA models tested whether the effects of glucocorticoids on risky habitat selection changed after calving and with increasing energy investment in lactation. Our model included distance to forest and shrubland (i.e., safer habitat) at the end of each step, an interaction between distance to forest and shrubland and FGM at the start of the movement bout, a three-way interaction between the distance to cover-FGM interaction and period before or after calving, and another three-way interaction between the distance to cover-FGM interaction and time since calving. We included movement bouts surrounding FGM samples collected up to 28 days before calving and up to 84 days after. To limit our inferences to locations associated with known FGM levels, we subsampled GPS data to the 20–h preceding each sample (i.e., within the period required for glucocorticoid metabolism). We sampled available steps from gamma distributions (turn angles) and von Mises distributions (step lengths) parameterized with movement characteristics of used steps (Avgar et al., 2016). We determined how many available steps were required to estimate selection coefficients by repeatedly fitting the model using ratios of between 1 and 1,000 available: used steps. Finally, to account for correlation between samples from individuals (Hebblewhite and Merrill, 2007), differences in sample size among individuals, and individual differences in habitat selection (Gillies et al., 2006), we included random intercepts for movement bouts, and random intercepts and slopes for all fixed effects and interactions. However, random effects models are challenging to fit within the conditional logistic regression framework typically used in step selection analysis because of the large number of step-specific strata. To deal with this challenge, we reformulated the conditional logistic model as a Poisson model with large, stratum-specific fixed intercepts (Muff et al., 2020) using the *glmmTMB* package in R (Brooks et al., 2017).

We used relative selection strength (RSS) a measure of habitat selection effect size. We calculated RSS across the 0.2–0.8 quantile range of FGM levels in the population (approximately 1,200–2,600 µg•g^-1^). RSS quantifies the ratio of the relative strength of selection for one location compared to selection at another location. When a single habitat characteristic varies between locations, RSS quantifies the change in selection for that characteristic (Avgar et al., 2017). In our case, we quantified the RSS for distance to cover habitat at the 0.2 quantile FGM versus a range of FGM values over the 0.2–0.8 quantile range. The difference in selection strength across this range predicts the change in effect size for selecting distances further from cover habitat as FGM increases. We compared the difference between these effect sizes by calving period while holding days since calving constant at zero, and differences by days since calving while holding calving period constant at post-calving.

We validated our model with used-habitat calibration (UHC) plots using the *uhcplots* package in R (Fieberg et al., 2018). UHC plots measure model calibration, or the agreement between distributions of habitat values at observed locations and distributions of habitat values at locations predicted as used by the model. UHC plots also compare used distributions to the distributions of habitat values at available locations to determine whether model covariates are important for predicting selection. Unlike other methods, UHC is appropriate for validating stratified habitat selection analyses like iSSA (Fieberg et al., 2018).

## Results

We used 68 fecal glucocorticoid metabolite samples from between May 14 and August 16 in 2019 and 2020, representative of 13 sampled individuals. Individuals each had between one and 16 fecal samples (median = 4) and between 14 and 554 location points (median = 153) across the pre- and post-parturition periods (Fig. S1). We used a ratio of 40 available: used points for all models as our sub-analysis suggested model coefficient estimates and standard errors remained relatively consistent from 30 to 100 available: used points (Fig. S2). Though individual sample sizes and location points per individual were few, small samples are still sufficient for RSF inference when selection strength is strong and landscape heterogeneity is low (Street et al. 2021).

We found no difference in glucocorticoid levels before and after calving (estimate = 0.10, 95% CrI −0.46, 0.65), though variation in glucocorticoid production was higher after calving. Figure 1). Despite no overall difference in production, selection for locations relative to forest and shrubland depended on glucocorticoid levels and changed over the calving season. In general, elk selected for locations closer to shrubland and forest (e^ß^ = 0.90, 95% CI 0.82, 0.99). They were 50% more likely to select locations further from shrubland and forest, and thus further into cropland, for each unit increase in glucocorticoid levels (e^ß^ = 1.44 95% CI 1.01, 2.04). This selection for cropland did not change immediately after calving when nutrition requirements were high (e^ß^ = 0.80, 95% CI 0.50, 1.28; Figure 2a). However, elk exhibited gradually stronger selection for locations further from forest and shrubland as days increased after calving (e^ß^ = 1.01, 95% CI 1.00, 1.02; Figure 2b). Our UHC model validation supported these inferences as model coefficients discriminated used from available locations. The model was well calibrated, with observed habitat use close to that predicted by models, and differences in the distribution of used and available locations with distance to shrubland and forest (Fig. S3).

**Figure 1.**
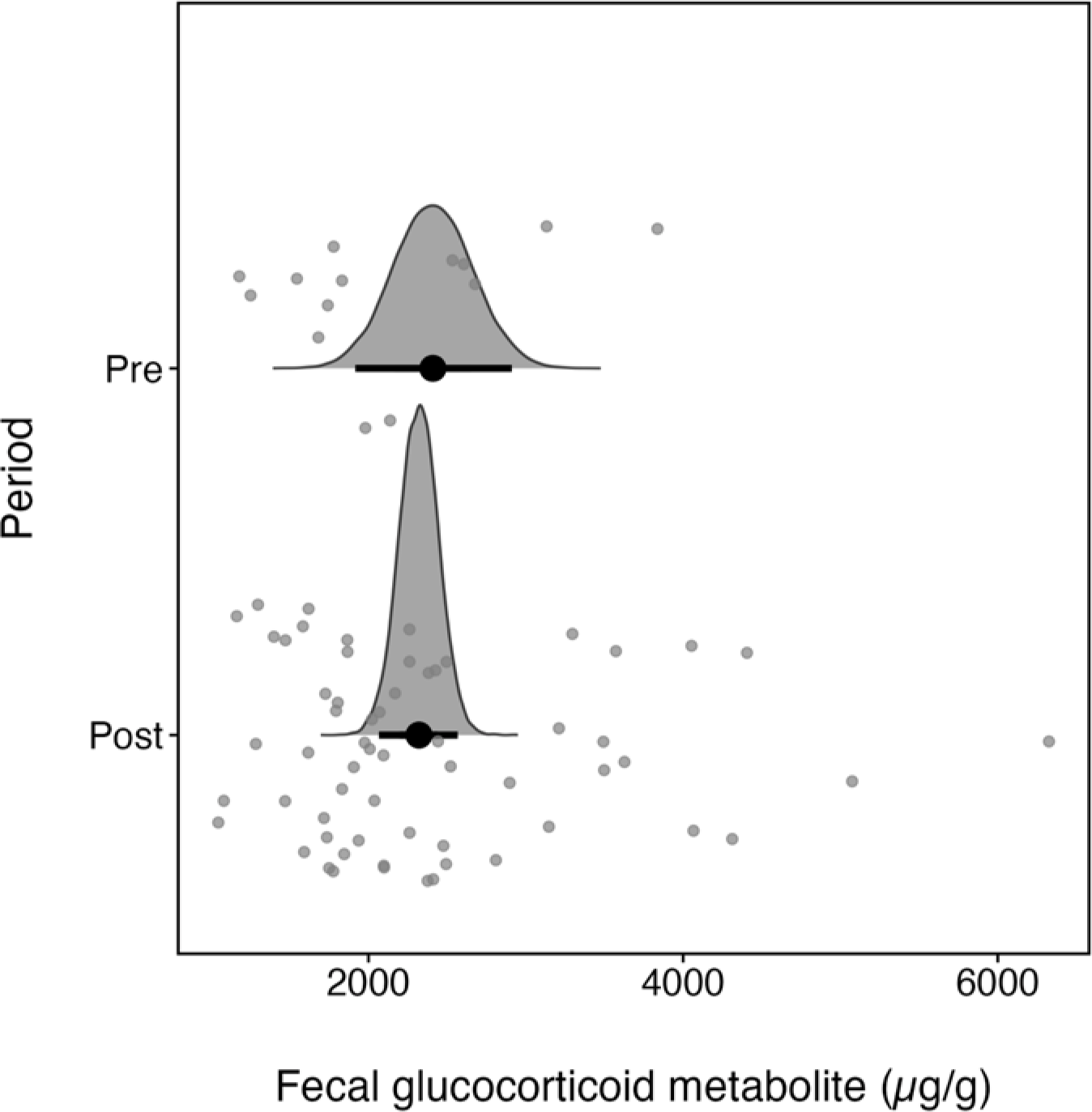
Half-eye plots comparing the distributions of glucocorticoid metabolite concentrations in fecal samples before (pre) and after (post) calving. Grey points are the glucocorticoid metabolite concentrations in each sample (n = 68), black points are the medians of the posterior distributions, and black intervals are the 95% quantile intervals of the posterior distributions.

**Figure 2.**
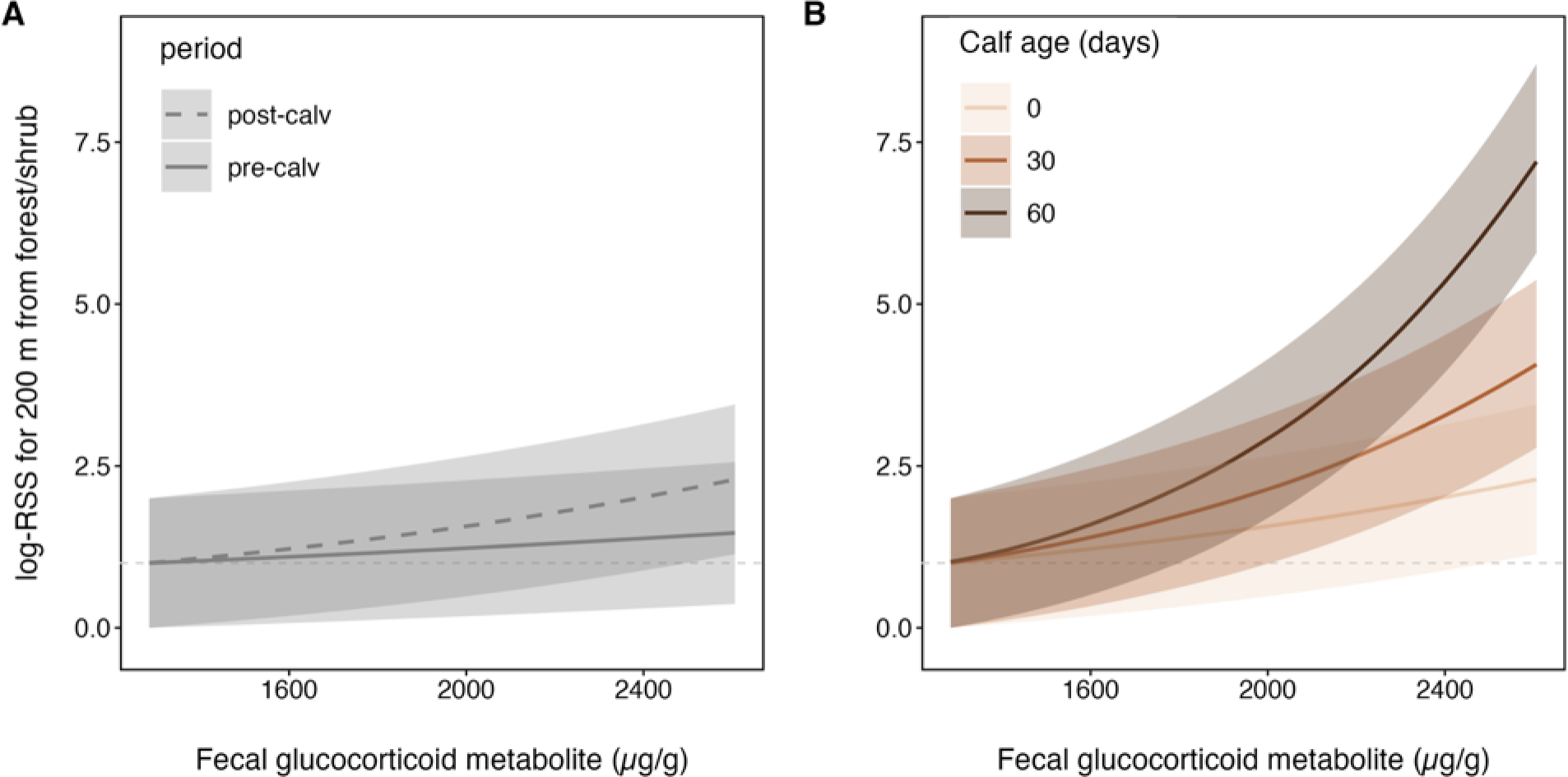
Log relative selection strength (RSS) for a location within forest and shrubland versus a location 200 m from cover with increasing glucocorticoid levels. In general, higher glucocorticoid levels predict greater selection for locations further from forest and shrubland, and thus closer to cropland. Panel A compares the RSS for locations relative to forest and shrubland before calving (pre-calv) and after calving (post-calv). Panel B compares RSS for these locations on the day of calving with calves aged 30 and 60 days. Solid lines are the mean predicted RSS and ribbons are 95% bootstrapped confidence intervals.

## Discussion

We used habitat selection models to test whether glucocorticoids drive the habitat selection behaviours that mediate trade-offs between parent and offspring survival. We found elk with elevated glucocorticoids selected more for cropland, which offers higher-quality forage, suggesting glucocorticoids drove them to accept more risk to meet their energetic needs. The cort-adaptation hypothesis theorizes glucocorticoids should encourage animals with offspring to take more risks to meet the demands of reproduction (Bonier et al., 2009a). We therefore predicted glucocorticoids might drive even stronger selection for risky foraging habitat after calving when selecting better forage would provide energy for elk to provision their calves. Contrary to our expectations, however, elk neither produced more glucocorticoids after calving, nor was there an abrupt change in the effect of glucocorticoids habitat selection after calving. On their own these results suggest glucocorticoids might not mediate trade-offs between parent and offspring survival in elk. However, the effect of glucocorticoids on cropland selection did increase with time since calving (Figure 2b), suggesting parents gradually accepted more risk as the energetic needs of calves grew. In the paragraphs to follow we discuss how this gradual rather than abrupt change in the effects of glucocorticoids on habitat selection provides a physiological mechanism for shifts in parental investment across life history stages.

We predicted an abrupt shift in the investment parents were willing to make in risk-taking behaviour after calving. We may not have found this abrupt shift because elk in our study system valued future reproductive opportunities over current offspring. Long-lived iteroparous mammals like elk have extremely high survival rates relative to their offspring, which makes any current reproductive investment relatively less valuable than the many more opportunities they will have to reproduce in the future (Gaillard et al., 1998). However, reproductive opportunities are fewer as aging animals approach reproductive senescence and current reproduction becomes more important for fitness. We had no age data for the adult female elk in our study system, but future studies might test whether glucocorticoids start to drive riskier habitat selection decisions more abruptly after calving for animals closer to reproductive senescence. Supporting the idea that glucocorticoids are related to shifts in parental investment, other studies have found glucocorticoid production changes with age (Heidinger et al., 2006; López-Jiménez et al., 2017). Either changes in glucocorticoid production or changing effects of glucocorticoids in older parents would follow theoretical expectations that individuals nearing senescence should invest more in offspring (Stearns, 1992).

Parental investment theory also posits that older offspring are more important to protect than younger offspring because parents have had more time to invest effort and energy into older offspring (Clark, 1994; Trivers, 1972). As predicted, we found glucocorticoids had a stronger effect on risky habitat selection by elk with older calves, with nearly three-fold stronger selection for the same glucocorticoid level when the calf reached 60 days compared to their day of birth (Figure 2). The energy invested in calves by 60 days would have presumably tripled their value to parents and warranted more risk-taking, including foraging in riskier but higher-quality cropland. Elk calves double in weight within the first 50 days (Nelson and Leege, 1982), suggesting the three-fold greater responsiveness to glucocorticoid production by female elk by 60 days might be proportional to or exceed their past investment. Empirical evidence from other species also supports the proportionally stronger investment parents are willing to make in older offspring as their energetic needs grow. For example, house wrens (*Troglodytes aedon*) invested more time alarm-calling in response to predators in proportion to the ages of their nestlings (Fernandez and Llambías, 2013). Our habitat selection models reveal glucocorticoids as a potential mechanism behind that proportional greater investment in risk for both elk and other species.

In addition to showing glucocorticoids are a mechanism for habitat selection, our study might also shed light on the physiological process that alters their effects after calving. In birds, behavioural investment in offspring frequently increases in proportion to circulating glucocorticoid levels (e.g., Bonier et al., 2009b; Bowers et al., 2019; Vitousek et al., 2014). Thus, changes in glucocorticoid production provide a physiological basis for parental behaviour. In contrast, many studies report no difference in glucocorticoid levels despite still finding changes in investment (e.g., Love et al., 2004; Patterson et al., 2011). In some of the latter studies, glucocorticoid effects on behaviour are thought to be mediated by circulating proteins called corticosteroid binding globulins (CBGs; Love et al., 2004). CBGs, which bind with high affinity to glucocorticoids in the bloodstream, regulate the “free” fraction of circulating glucocorticoids able to bind to receptors in the body (Breuner et al., 2020). Some of these glucocorticoid-activated receptors appear to regulate offspring care behaviour, making the free fraction most important for behaviour. For example, elevated free but not total glucocorticoid levels caused European starlings (*Sturnus vulgaris*) to abandon their nests (Love et al., 2004). Despite finding that glucocorticoids in our study increased investment in risky habitat selection as offspring age, we similarly found no difference in total glucocorticoid levels after calving. It is possible risky habitat selection gradually increased after calving not because glucocorticoid production gradually increased, but because CBG levels gradually declined. Integrating CBGs into habitat selection models could help test whether they help mediate investment between parent and offspring.

Two factors unrelated to the value of the calf might have also influenced the gradually changing effects of glucocorticoids on habitat selection by female elk after calving. First, glucocorticoids might have had a gradually stronger effect on selection for riskier habitats after calving because older calves require less care. Elk, like most other mammals, remain with their parents for an extended period of offspring care after birth. In addition to dramatically increasing the energetic needs of female elk during this period, elk calves have limited mobility for up to three weeks after birth (Geist, 1982). This initial period of limited mobility in our study likely also limited movement by female elk, and thus the range of risky habitats they could access, with the effects gradually relaxing as calves became more mobile. Birds also care for their offspring over an extended period after birth. However, many nestling birds receive biparental care and remain in a central nest location, freeing at least one parent to move in search of forage quality. Repeating our habitat selection study using bird species or other animals whose offspring do not constrain their movements might isolate the effect of offspring energetic needs on glucocorticoid-driven habitat selection by parents.

It is also possible elk gradually used riskier habitat after calving because of a gradual improvement in forage quality. If forage quality matured alongside calves, it might have been worthwhile for female elk to accept more risk while foraging. Risk-taking for higher-quality forage is a common strategy among reproductive animals supporting offspring. The reproductive phenologies of birds, mammals, fish, and many other species, for example, have adapted to coincide with risky, long-distance migrations to arrive on summer ranges when their offspring will benefit most from forage quality (Alerstam et al., 2003). Elk similarly time offspring production to peak forage quality, with calving generally beginning mid-May and ending mid-June to coincide with early plant growth (Skovlin, 1982). However, plant growth is an unlikely explanation for the stronger effects of glucocorticoids on risky habitat selection in our study system because crop seeding dates vary, meaning some crops were likely in their early nutritious stages throughout or even before the calving season. Given the variation in timing of crop maturity but consistent habitat selection behaviour by elk after calving, we contend the most likely explanation for the gradually stronger effect of glucocorticoids on cropland habitat selection remains the energetic needs of growing calves.

By foraging in cropland to support their calves, elk likely also encountered more stressors, which may have encouraged them to select even more strongly for cropland. Activation of stress responses release glucocorticoids that stimulate feeding behaviour (White et al., 1994) independent of the energetic needs of calves. We did find potential evidence for more stress responses after calving. While we found no significant differences in glucocorticoid production before and after calving, there was more variation in glucocorticoid production after calving, including higher peaks (Figure 1). The need to forage in risky cropland might initially been caused by the energetic needs of calves, but it is possible spending more time in cropland created a feedback loop between glucocorticoid production and habitat selection that caused elk to continue to select for cropland. Thus, while energetic needs were likely the primary driver behind the effects of glucocorticoids on habitat selection, the gradual change might have also been influenced by habitat selection itself. Since theory suggests habitat with the most forage is also often the riskiest in terms of predation (Brown and Kotler, 2004), parents of many species probably face similar feedback loops as their offspring age.

In our study, we demonstrated glucocorticoids drive habitat selection behaviour. While glucocorticoids are known to affect many behaviours involved in reproduction, their effect on habitat selection is promising for understanding the universal importance of glucocorticoids in reproduction because virtually all mobile animals must forage in risky habitats to provision offspring. Glucocorticoids have always been measured as a response to stressors, but testing how glucocorticoids affect habitation selection fosters thinking about how their effects change across life history stages. We suggest integrating glucocorticoid-integrated habitat selection models can help elucidate the fitness-enhancing role of glucocorticoids in stressful environments.

