## Supplementary Figs 1-3 for "Life history trade-offs in glucocorticoid-mediated habitat selection"

**Supplementary materials for *Life history trade-offs in glucocorticoid-mediated habitat selection***

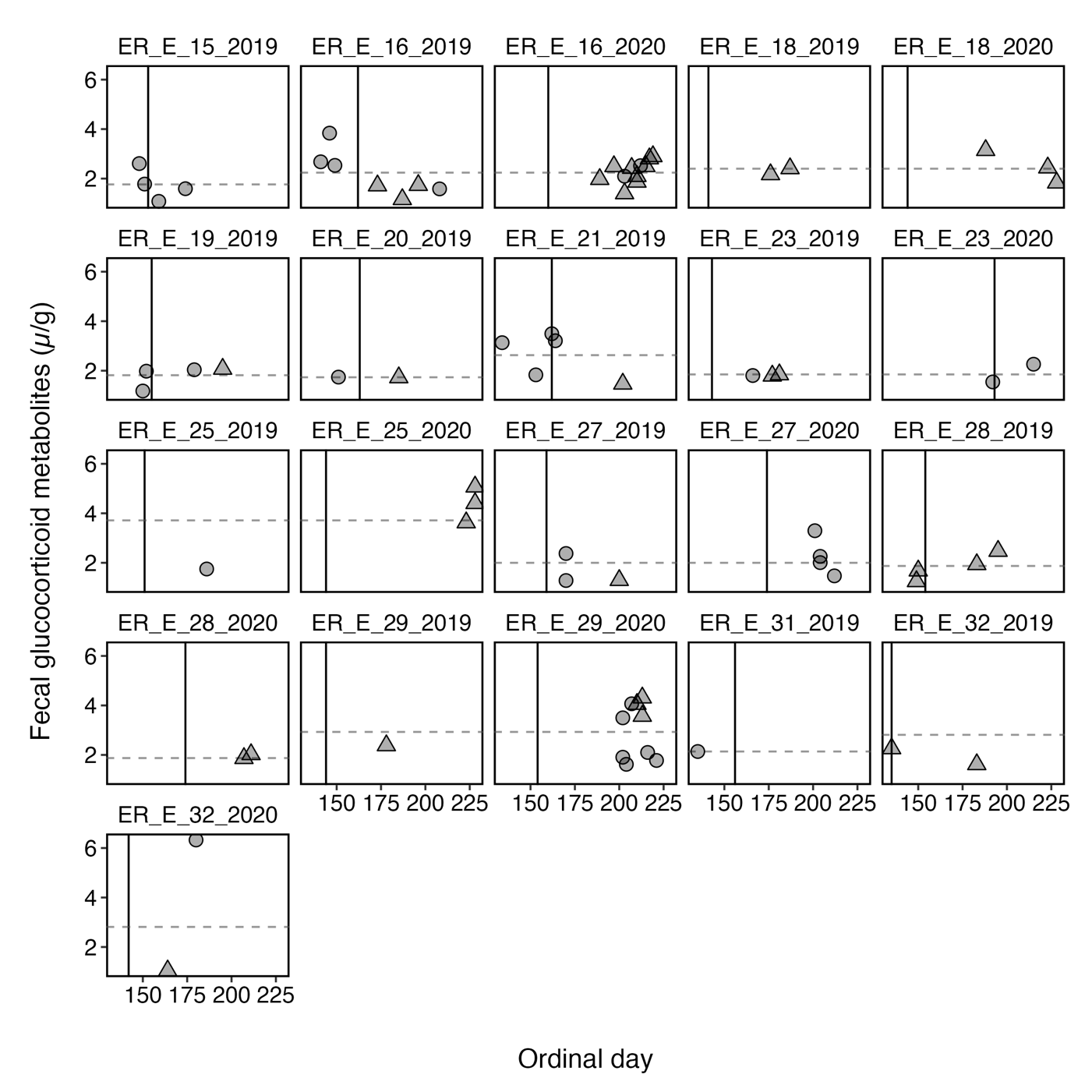
**Figure S1.** Collection dates and glucocorticoid metabolite levels from 68 fecal samples from 13 individual elk in 2019 and 2020. Each panel represents a single elk, with the vertical black line indicating the individual’s calving date for that year. Circles denote samples identified as belonging to individuals by matching DNA, and triangles denote samples identified as belonging to individuals based on machine learning. Horizontal dashed lines show the median fecal glucocorticoid metabolite level for individuals across both calving seasons.

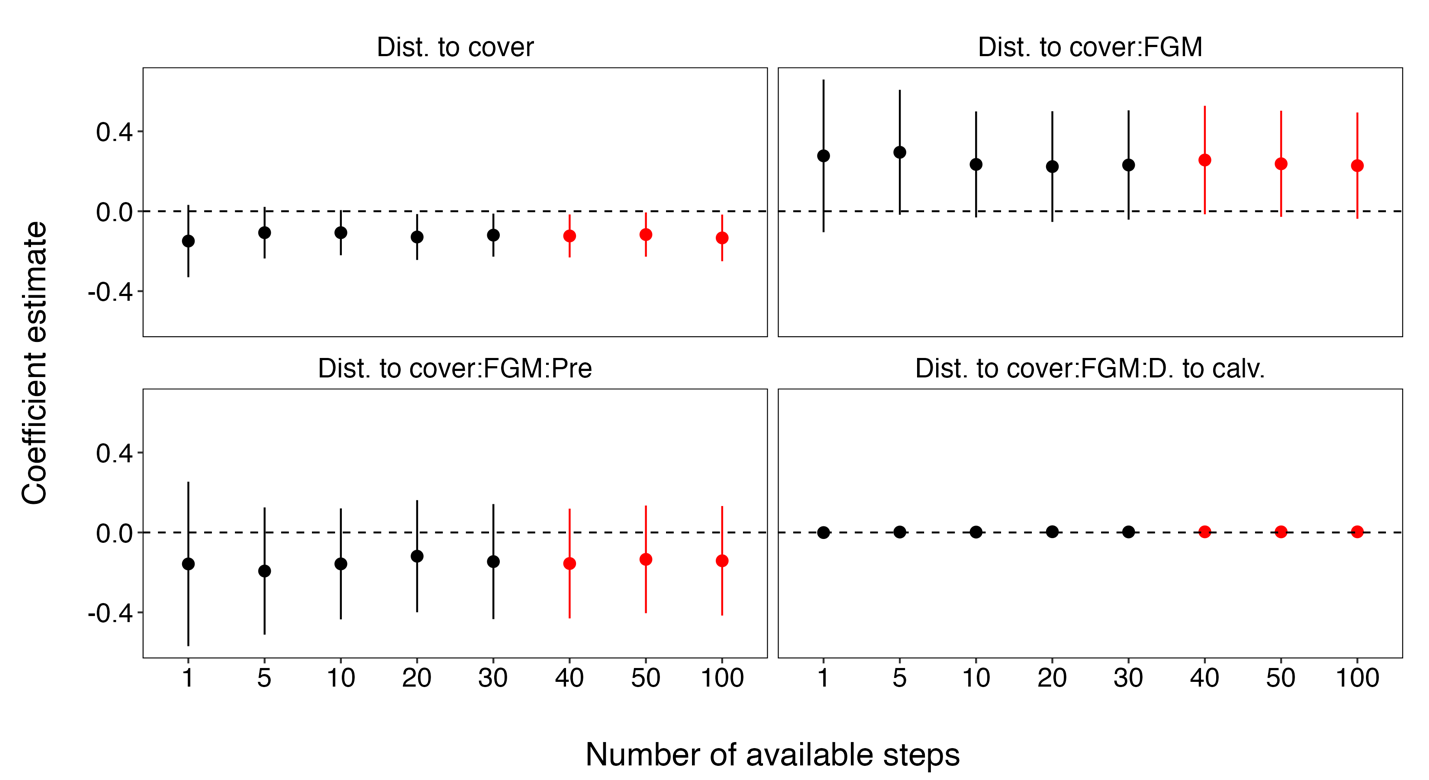
**Figure S2.** Model coefficient estimates (points) and standard errors (vertical lines) from fitted integrated step selection model across increasing ratios of available: used points. Black points and lines indicate available: used ratios below 40, and red points and indicate ratios above. Coefficients and standard errors remain relatively consistent at available: used ratio equal to 30. Pre = pre-calving period (reference category relative to post-calving period), and FGM = fecal glucocorticoid metabolite levels.

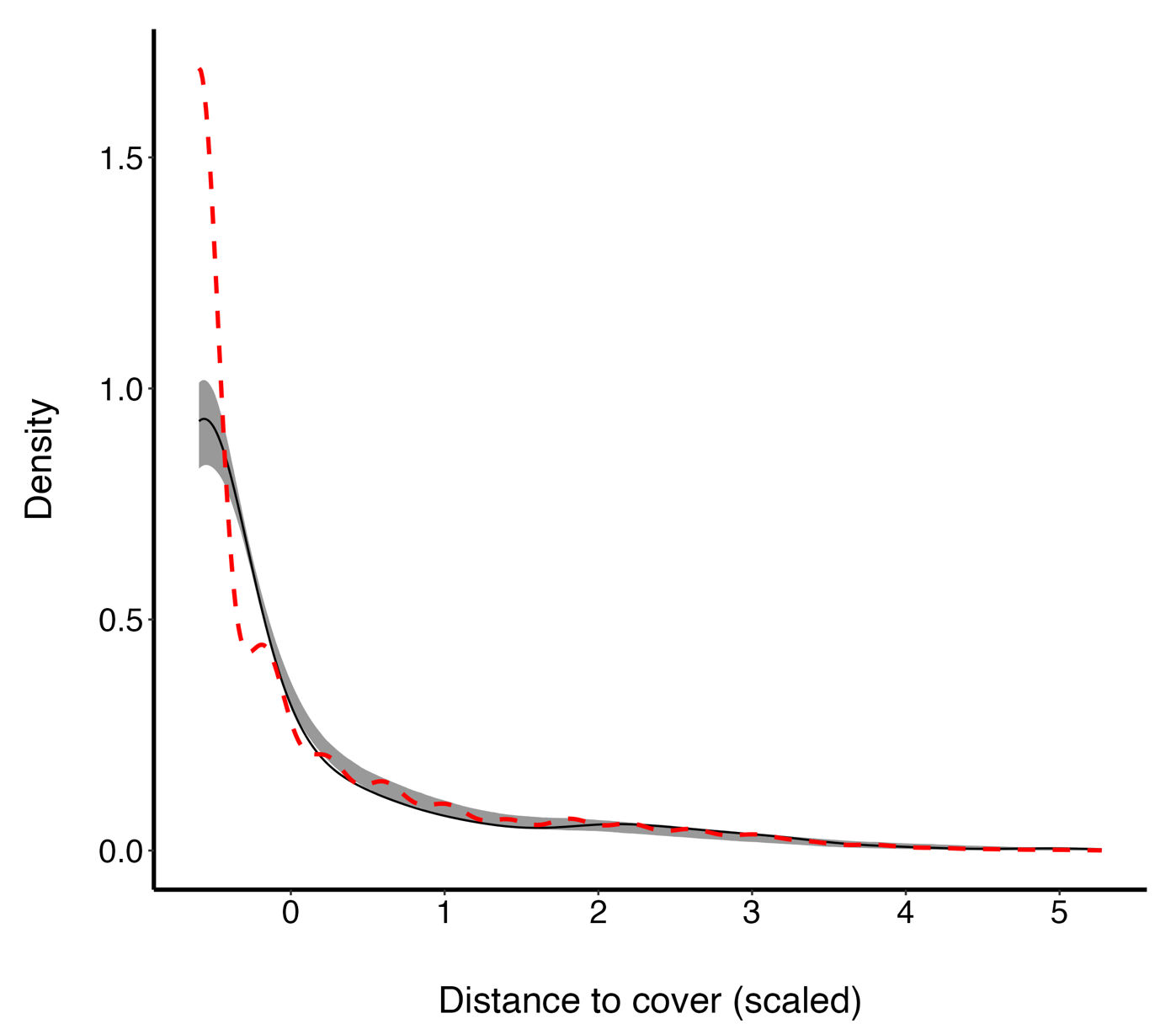
**Figure S3.** Used-habitat calibration plots comparing distributions of habitat values at used locations (black solid line), habitat values at available locations (red dashed line), and 95% simulation envelopes for the predicted distribution of habitat covariates expected at the used locations based on our fitted integrated step selection model (grey ribbon). Predicted values agree with observed values — i.e., the model is well-calibrated — when the model predictions (grey ribbons) overlap with observed values (black lines). Differences between the black-solid and red-dashed lines indicate elk are likely to be found at locations with dissimilar characteristics to those available.
